# reComBat-seq: Regularized negative binomial regression for batch-effect correction in underdetermined transcriptomics datasets

**DOI:** 10.64898/2026.05.27.728166

**Authors:** Zhasmina Stoyanova, Jörg Menche, Daniel Malzl

## Abstract

**Motivation:** Batch effect correction is essential for the integration of large-scale transcriptomics datasets such as single-cell RNA-seq or multi-study bulk RNA-seq datasets for reducing technical noise that may mask biological signal. Existing correction methods, either do not produce count data output which is crucial for state-of-the-art downstream analyses such as differential expression analysis or fail to converge in underdetermined study designs.

**Results:** We present reComBat-seq, a method that extends the Negative Binomial regression framework of ComBat-seq by incorporating Elastic Net regularization. This approach resolves problems with rank-deficient design matrices while also preserving the integer nature of count data. Benchmarking on simulated and real datasets such as single-cell RNA-seq data demonstrates that reComBat-seq successfully removes batch effects in complex study designs while maintaining compatibility with downstream differential expression tools.

**Availability and Implementation:** reComBat-seq source code can be found at https://github.com/menchelab/reComBat-seq. All code to reproduce the presented analyses can be found at https://github.com/menchelab/reComBatseq_Studies. Data produced in this study is available at https://doi.org/10.5281/zenodo.19736515. Used single-cell RNA-seq data can be found at https://doi.org/10.5281/zenodo.14234956.

**Supplementary Information:** Proofs and volcano plots of differential expression analysis

## Introduction

Modern analysis of biological data requires handling large datasets which can contain a substantial amount of noise. A specifically unwanted part of this noise is the batch effect which is introduced by technical differences in sample handling and data processing and often masks true biological signals. Thus removal of this technical variation is crucial for analysis of datasets composed of different published studies or even data generated on different days (Hui, *et al*., 2024; Goh, *et al*., 2022, 2017; Leek, *et al*., 2010).

A variety of approaches have been developed for the removal of such batch effects (Zhang, *et al*., 2020; Adamer, *et al*., 2022; Zhang, 2025; Lopez, *et al*., 2018; Korsunsky, *et al*., 2019; Johnson, *et al*., 2007) (see (Yu, *et al*., 2024) for a more comprehensive overview) with linear regression-based methods counting as some of the simplest and best understood ones. Some of the most widely used ones falling in this category are the methods of the ComBat family. Despite their frequent and successful use on RNA sequencing (RNA-seq) based data (Luecken and Theis, 2019; Adamer, *et al*., 2022; Wolf, *et al*., 2018), they were originally developed for microarray data. Their application thus requires substantial data transformation to make the negative binomially distributed count data from RNA-seq amenable for the method’s assumption that the data is normally distributed. This transformation not only makes the method prone to erroneous results but also hinders typical downstream analyses employed with RNA-seq data, such as differential expression analysis with tools like DEseq2 or edgeR (Love, *et al*., 2014; Chen, *et al*., 2025) which use count data as input. Furthermore, ComBat suffers from low performance in low sample high batch covariate designs in which the design matrix may be rank deficient and the method plainly fails due invertibility issues during fitting. Both of these points were recently addressed by the introduction of ComBat-seq (Zhang, *et al*., 2020) and reComBat (Adamer, *et al*., 2022), where the former now respects the count nature of sequencing-based data and the latter introduces regularization to ensure invertibility of the design matrix. However, current datasets not only primarily consist of sequencing data but are increasingly complex with respect to experimental design. Thus, to effectively reduce or remove batch effects, dedicated methods need to solve both aforementioned issues.

To address this, we developed reComBat-seq. Much like reComBat, reComBat-seq extends the ComBat-seq method with elastic net regularization. This enables us to fit the model on rank deficient experimental designs while respecting the count nature of sequencing based data. We further show that reComBat-seq effectively integrates in regular RNA-seq analysis pipelines by comparing it against its predecessors with respect to differential expression analyses.

## Methods

### Elastic net expansion of ComBat-seq

ComBat-seq aims to estimate the parameters of a negative binomial distribution (*Y*_*ijg*_ ~*NB*(μ_*ijg*_, ϕ_*ig*_) for each gene *g* with respect to sample *j* and batch *i* by solving the equation:

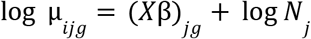

where μ_*ijg*_ is the expected value for the negative binomial of distribution of read counts for gene *g* in sample *j* and batch *i, N*_*j*_ is the sum of all counts in sample *j* (i.e. the library size), β is a vector of regression coefficients and *X* being the design matrix. The accompanying dispersion parameter ϕ_*ig*_ is computed using the Cox-Reid adjusted profile likelihood method (APL)

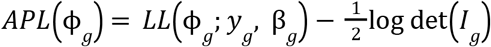

with *LL*(ϕ_*g*_; *y*_*g*_, β_*g*_) being the log-likelihood of ϕ_*g*_ given the genes observed expression *y*_*g*_ and estimated regression coefficients β_*g*_ and *I*_*g*_ being the Fisher information matrix. ComBat-seq solves these equations via maximum likelihood estimation of β by finding the minimum of the negative log-likelihood function

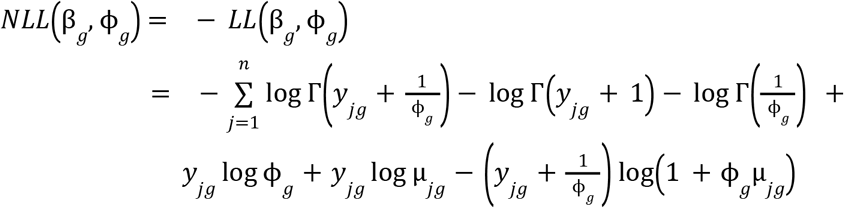

This is done by iteratively solving

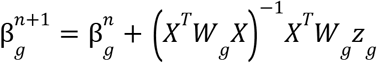

until convergence. Here, *X*^*T/*^*W*_*g*_ *X* is equivalent to the Fisher information matrix *I*_*g*_, where *W*_*g*_ is a diagonal weight matrix containing the expected second derivatives of the negative log-likelihood function and *z*_*g*_ is the working response, i.e., the linearized version of the observed counts around the current parameter estimate that allows the non-linear problem to be solved via weighted least squares (see Supplementary information for full derivation). This procedure relies on the invertibility of *I*_*g*_ and thus on the design matrix *X* being full rank. Thus, ComBat-seq fails to fit on datasets with rank-deficient design matrices such as those that occur in meta-analysis settings like integrating datasets from multiple studies where-dimensional covariates may be linearly dependent. To alleviate this, we reformulated ComBat-seq in a regularization framework. Specifically, we introduce elastic net in the estimation of the regression coefficients which lets us rewrite the target function using *NLL*(β_*g*_, ϕ_*g*_) from above as follows

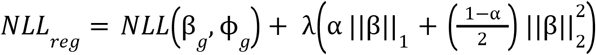

where λ determines the strength and α the type of regularization with α = 0 corresponding to pure LASSO and α = 1 corresponding to pure ridge regression. 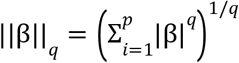 is the *l*_*q*_ norm for *q* ≥ 1 with *p* being the number of covariates. This lets us rewrite the iterative update step in IRLS as

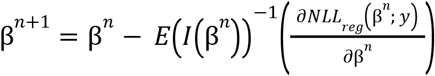

where *E*(*I*(β)) is the expected Fisher information matrix computed as

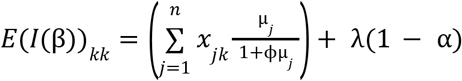

and the gradient of *NLL*_*reg*_ as

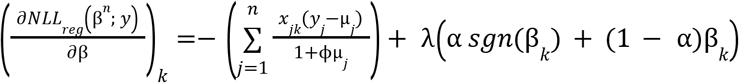

with respect to the *k*-th coefficient, β_*k*_.

In brief, incorporating the second derivative of the ridge penalty into the diagonal of the Fisher information matrix ensures invertibility in rank-deficiency settings. See supplementary information for full derivation of the above.

### Implementation of reComBat-seq

reComBat-seq is implemented in R and builds on code from the ComBat-seq (Zhang, *et al*., 2020) and edgeR (Chen, *et al*., 2025). To incorporate regularization, reComBat-seq modifies the core fitting function by extending the objective function to include the elastic-net penalties. Specifically, the regularized version alters both, the gradient by adding the first derivatives of the penalty terms to it and the Fisher information matrix by incorporating the second derivatives into its diagonal. These changes enable elastic net–style penalization while maintaining the structure of the IRLS-based optimization routine. The specific routines are as follows

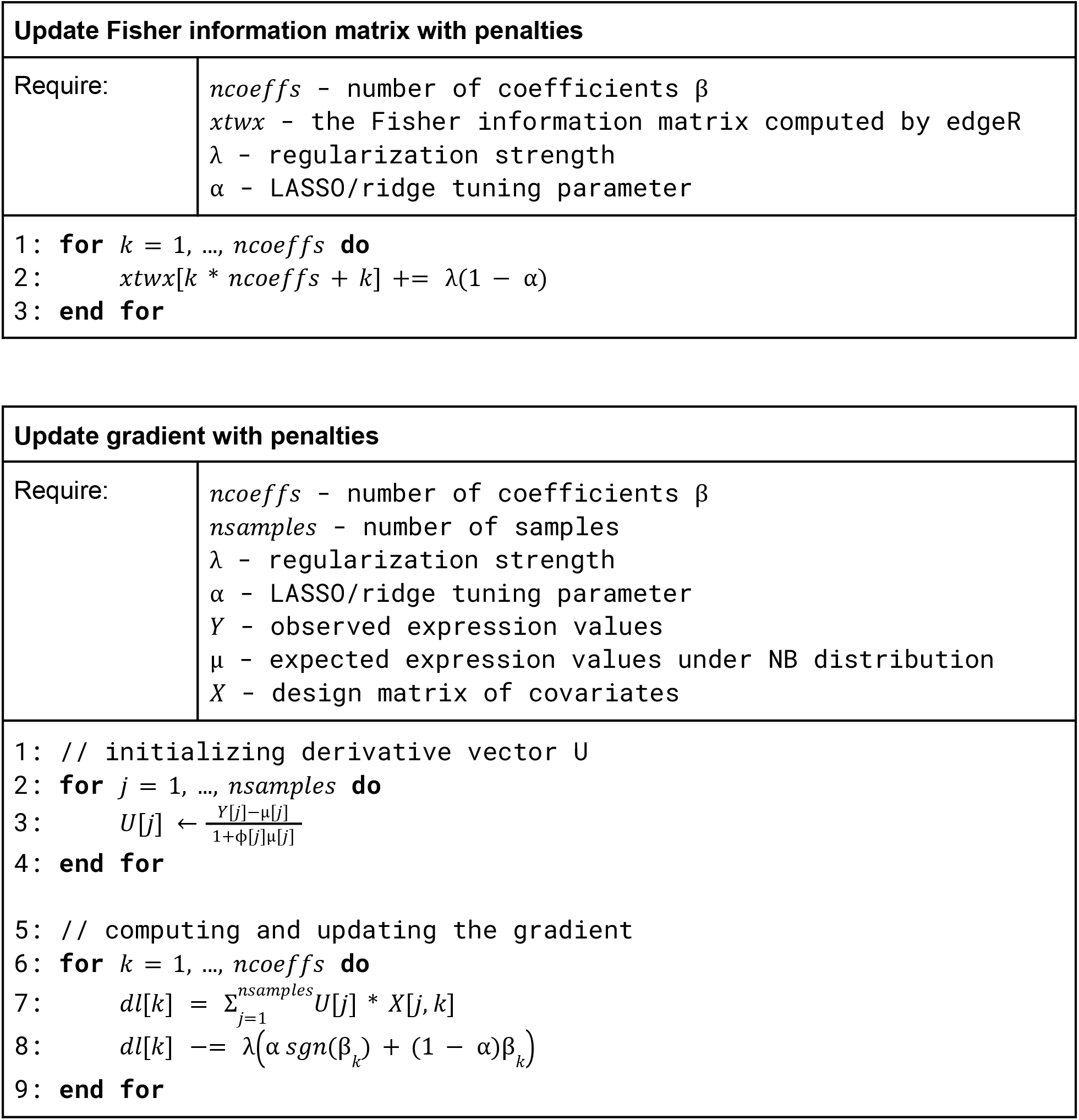

### Parallelisation

Since the model is fit for each gene independently, the framework lends itself naturally to parallelisation. We employ user-configurable multithreading via OpenMP to distribute the gene-level parameter estimation across available CPU cores. To ensure memory efficiency and thread safety, each thread operates within a private workspace containing local buffers for offsets, dispersions and weights. The parallel architecture effectively offsets the complexity of the regularization framework. Performance tests indicate that with a minimal allocation of two threads, the execution time of reComBat-seq remains on par with the original ComBat-seq implementation.

### Performance evaluation of reComBat-seq

To evaluate the performance of the presented tool we compared reComBat-seq to its predecessors ComBat, ComBat-seq and reComBat (Adamer, *et al*., 2022; Zhang, *et al*., 2020; Johnson, *et al*., 2007) in a series of simulation experiments. Borrowing from the design used in Zhang et al. 2020 and Tran et al. 2020 (Zhang, *et al*., 2020; Tran, *et al*., 2020) we built a pipeline that consists of (i) data simulation using Splatter (Zappia, *et al*., 2017), (ii) adjusting the introduced batch effects using the dedicated tools and (iii) benchmarking the results visually using principal component analysis (PCA) and uniform manifold approximation and projection (UMAP) as well as via quantitative evaluations like examining the true positive and false positive rate (TPR and FPR) and precision in terms of differentially expressed genes compared to the simulated ground truth.

### Data simulation

To test our algorithm against a known ground truth we simulated datasets containing batch effects. In brief, we used splatter to generate synthetic RNA-seq datasets across different parameters. Four main experiments were designed to evaluate reComBat-seq under controlled conditions and compare it to the aforementioned methods. An initial hyperparameter study examined the impact of regularization tuning on the overall performance. As no consistent trend was observed, the rest of the experiments utilized a fixed tuning parameter (α = 0.3) and regularization strength (λ = 0.8). See Figure S1 for the full results of the hyperparameter screen. Subsequent experiments isolated the impact of key dataset characteristics: (i) number of genes, (ii) samples, and (iii) batches. In each case, one variable was varied while the others were maintained at a fixed baseline of 4,096 (for genes/samples), and two (for batches). Initial parameters were estimated from the airway (Himes, *et al*., 2014) dataset, with batch effect factors drawn from normal distribution (batch.facLoc ~ *N*(2, 0. 3), batch.facScale ~ *N*(0. 8, 0. 15)). In all scenarios, we simulated two groups with 10% of genes designated as differentially expressed (de.facLoc = 0.5, de.facScale = 0.2). Ground truth was established by re-simulating the biological signal without batch effects (batch.rmEffect = TRUE). Furthermore we introduced a confounded design matrix to evaluate the performance in rank-deficient settings.

### RNA-seq data

Comparison of reComBat and reComBat-seq was also carried out on real world data. In brief, we downloaded sequencing data from several studies of psoriasis (SRP238713, SRP065812, SRP035988, SRP026042, ERP110816) from the Sequencing Read Archive (SRA) using the nf-core fetchngs pipeline v1.7 (Patel, Beber,, *et al*., 2022). We then uniformly processed the data with the nf-core rnaseq pipeline v3.7 (Patel, Ewels,, *et al*., 2022) aligning against the GENCODE human genome assembly release 40 with STAR. We then used subread’s featureCounts to count reads per gene against the accompanying comprehensive gene annotation from GENCODE allowing only non-duplicated, single overlap, exon-mapping reads to be summarized by gene id (-t exon -g gene_id -T 10 --ignoreDup). The resulting count matrix was then subjected to batch correction using either reComBat v0.1.4 or reComBat-seq using the SRA study accession as batch and retaining disease annotation as wanted variation. To adhere to reComBat’s assumptions about the data distribution we applied log + 1 counts per million normalization and subsequent row-wise standardization before running reComBat. Data was supplied as is to reComBat-seq.

### Differential expression analysis

We performed differential expression analysis for all obtained results including individual experiments. For raw and reComBat-seq corrected data, we applied the standard DESeq2 (Love, *et al*., 2014) pipeline including shrinkage for raw data. For reComBat corrected data we used the standard limma (Ritchie, *et al*., 2015) pipeline. Differential gene expression for scRNA-seq data was conducted as described in (Neuwirth, *et al*., 2025). In brief, we used MAST v1.36.0 (Finak, *et al*., 2015) with prior normalization to counts per 10,000 UMIs. Genes were found significant if (i) p_adj_ < 0.001 and abs(log2FC) > 0.5.

### Differentially expressed genes recall

To assess recall of differentially expressed genes (DEGs) after batch corrections, we compared DEGs computed from the full and individual datasets to those found in the uncorrected data. For this, we computed DEGs for all raw datasets individually and aggregated the top and bottom 300 DEGs with p_adj_ < 0.05 into two ground truth sets by either a simple union (union recall) or by only adding those that are found in at least three of the five studies considered (intersection recall). We then computed the recall as the size of overlap of DEGs of the full, corrected datasets with the ground truths in percent. For individual dataset recall, we computed DEGs on the individual corrected datasets (i.e. correcting the full dataset and then splitting it again) and compared the top and bottom 300 DEGs to the raw dataset DEGs for different sets of cutoffs. Here we chose to use the log2-fold change (LFC) cutoff divided by three as the cutoff for reComBat corrected data as they tended to be only a third of the raw / reComBat-seq log2-fold changes on average. Chosen cutoffs are the α ∈ {0. 01, 0. 05, 0. 1} and *LFC* ∈ {0. 25, 0. 5, 1}

### Single-cell RNA-seq data

We used a previously published scRNA-seq dataset from (Neuwirth, *et al*., 2025). In brief, we downloaded the data from Zenodo (Malzl and Neuwirth, 2024) and used the raw count data contained within the ‘raw’ member of the loaded AnnData object.

## Results

Ease of use and relatively low costs have not only made gene expression analysis via RNA sequencing an indispensable tool for biological research but also resulted in the development of a myriad of different isolation protocols, preparation techniques and sequencing methods. These not only differ in the way they isolate, prepare and sequence RNA from a cell or a tissue but also tend to introduce technical variation due to differences in isolated RNA content, sequence recovery by the instrument and even the experimenter handling the samples. Removing this non-biological noise is vital for the analysis of datasets consisting of multiple experiments as it can mask biological signals and reduce statistical power (Leek, *et al*., 2010; Goh, *et al*., 2022, 2017). Similar to RNA-seq methodologies, there have been many attempts to counteract this nuisance variation, but despite their success most of these methods assume Gaussian distributed data. This assumption not only violates the overdispersion seen in count-based data but also fails to provide results consistent with other count-native analysis tools like DESeq2 or edgeR (Zhang, *et al*., 2020; Anders and Huber, 2010). Those methods that do are incapable of handling the complex designs demanded by modern investigations. To change this we developed a tool which combines the count-nativity of ComBat-seq (Zhang, *et al*., 2020) and the flexibility of reComBat (Adamer, *et al*., 2022) to handle large, complex datasets.

### reComBat-seq reduces false positives in differential expression analysis

The first step towards this goal was the adaptation of the edgeR framework to incorporate regularization. We used edgeR because of its established role in what we wanted to achieve as well as its efficient code. Once amended and tested we sought to compare reComBat-seq to its predecessor ComBat-seq and reComBat. We first simulated differential expression experiments for a range of number of batches, samples and genes using splatter. We then used the aforementioned tools to remove the batch effect from the data and subsequently performed differential expression analysis to assess how well each correction recovers the differentially expressed genes. Figure 1 shows the true and false positive rate (TPR and FPR) as well as the precision for all three tools (note that ComBat-seq is not shown in the variable batch experiments as it is not able to handle singular matrices). On the top level, all three perform equally well with respect to the number of recovered true differentially expressed genes with reComBat-seq seemingly dealing better with a larger number of genes in the dataset (Figure 1 top row). Despite reComBat approaching the count-based methods with increasing data amount, it clearly shows an elevated false positive rate compared to the latter (Figure 1 middle row). This indicates that the transformation necessary to comply with reComBats distribution assumption introduces artificial differences in the data which is also echoed by the precision (Figure 1 bottom row). Taken together, these results show that the performance of reComBat-seq is comparable to the state-of-the-art methods and even outperforms reComBat when it comes to avoiding spurious results after batch correction.

**Figure 1:**
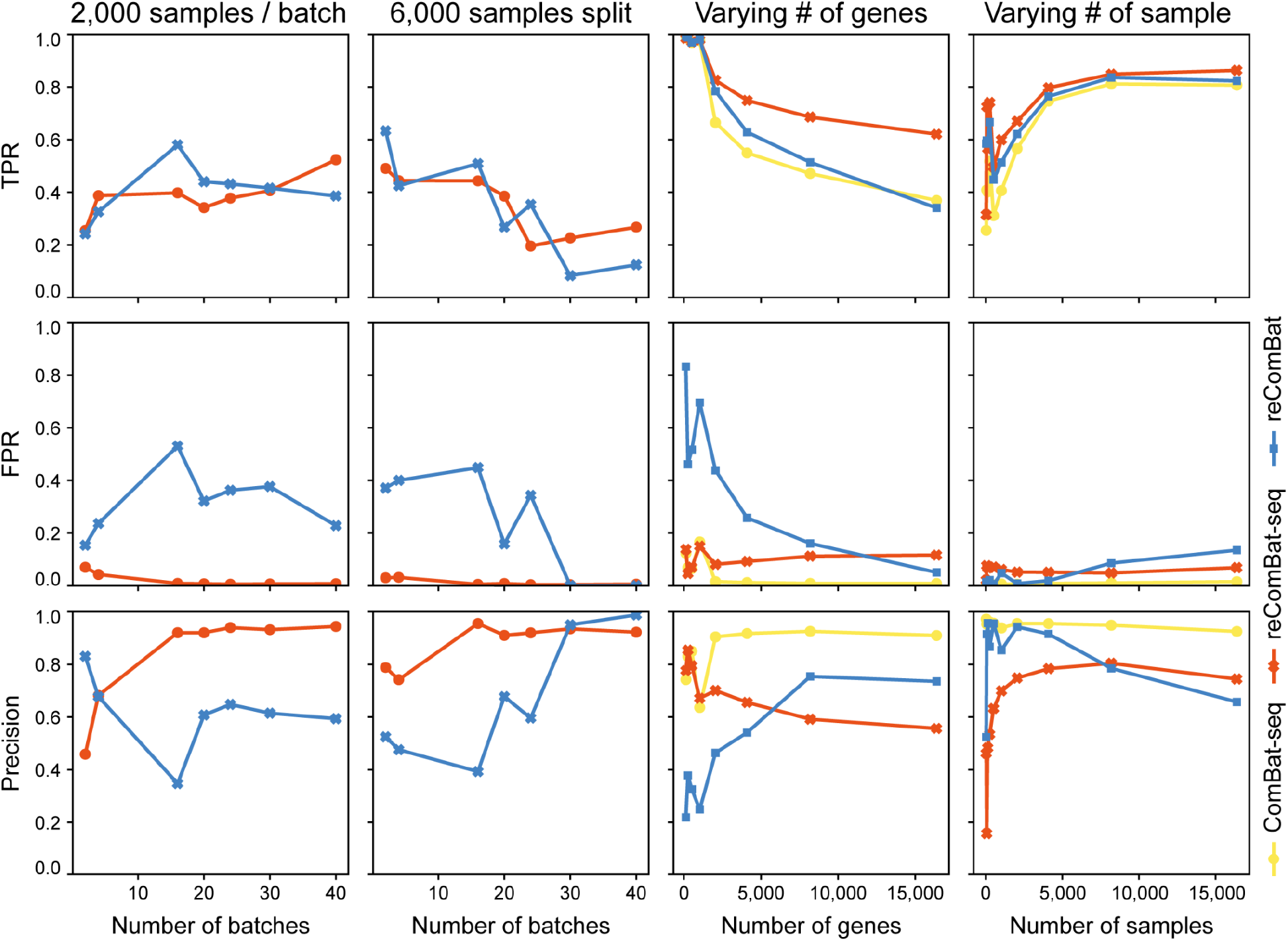
True positive rate (TPR), false positive rate (FPR) and precision plots over different simulation experiments. All values are computed as proxies of recall of the differentially expressed genes from simulated ground truth. Missing method in a plot means that the method could not be applied to the simulated data with singular design matrices.

### reComBat-seq effectively removes unwanted variation while retaining biological information

Simulation experiments are vital to benchmark methods against each other but they often do not reflect the behaviour against real world data. This prompted us to replicate the results obtained on simulated data with a real biological dataset. To this end, we used data of psoriasis patients and healthy controls from multiple studies. In total we downloaded 321 primary tissue samples of skin from five studies (Foulkes, *et al*., 2019; Li, *et al*., 2014; Di Meglio, *et al*., 2014; Gupta, *et al*., 2016; Yu, *et al*., 2020). 141 of them were from healthy volunteers and 180 came from psoriasis patients. After uniform processing from raw data to remove any variation introduced by differences in computational tooling, we performed batch correction with reComBat and reComBat-seq and assessed the effectiveness by PCA. Here both algorithms perform similarly well by effectively merging the different batch clusters to two distinct health and disease clusters (Figure 2A). Next we performed differential expression analysis on the corrected, individual uncorrected and individual corrected datasets. We used the results of the individual uncorrected as a benchmark for the corrected ones as proposed in (Hui, *et al*., 2024). In brief, we generated two ground truth gene sets, one based on the union and one based on the intersection of differentially expressed genes (DEGs). We did this by computing the top and bottom 300 DEGs of the individual uncorrected datasets with p_adj_ < 0.05 and then only retained those found in at least three of the five studies for the intersection-based and a simple union of all for the union-based set. We then considered the overlap in percent with respect to the ground truth sets as our recall metric. In total we found 309 DEGs in the intersection ground truth set of which reComBat recovered 262 and reComBat-seq recovered 296. This makes reComBat-seq slightly better at recall with 0.96 compared to 0.85 with reComBat. We saw a similar picture for the union ground truth set, where we found 1,863 DEGs of which reComBat recovered 1,009 or 54% and reComBat-seq recovered 1,122 or 60%. However, we found that reComBat corrected datasets have consistently higher adjusted p-values as well as consistently lower log-fold changes as compared to uncorrected and reComBat-seq corrected datasets (Figure S2). This raised the question of how well the individual corrected datasets recover the differentially expressed genes of the individual uncorrected ones, since the ultimate goal of batch correction is to remove technical variation while keeping biological variation possibly untouched. We thus computed DEGs similar to before finding the top and bottom 300 DEGs after significance filtering and calculated the recall for each dataset individually. Finally, we did this for a range of commonly used significance thresholds to assess the stability of the results with respect to the significance filtering. Despite the overall comparable results for the full corrected datasets, the individual comparisons show that reComBat-seq retains more of the original biological information over all used thresholds (Figure 2B). Interestingly, the regularization of the batch correction also has a variance stabilizing effect on the data leading to comparable results with and without log-fold change shrinkage. In summary, reComBat-seq outperforms reComBat when it comes to removing unwanted variation while retaining biological information contained in the datasets.

**Figure 2:**
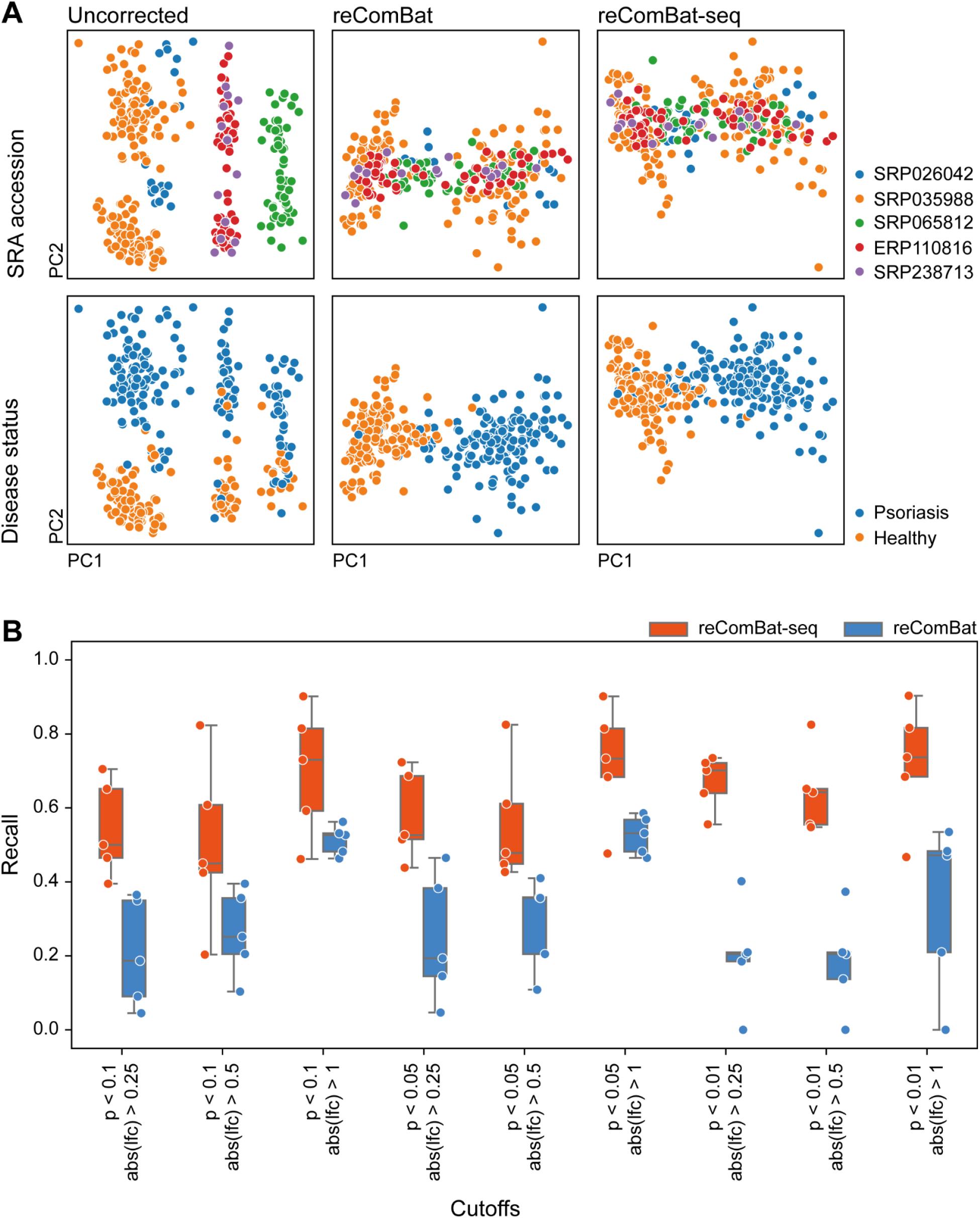
Comparison of reComBat and reComBat-seq on real data. (A) PCA scatter plots of uncorrected, reComBat and reComBat-seq corrected data. Uncorrected and reComBat-seq corrected data was log(cpm + 1) transformed before PCA. (B) Recall of differentially expressed genes for individual datasets after correction with respect to uncorrected data.

### reComBat-seq can be readily applied to single-cell RNA-seq data

After finding reComBat-seq’s performance being on par with reComBat and since its predecessor ComBat is still used in the popular scanpy framework (Wolf, *et al*., 2018), we asked whether we could use reComBat-seq to integrate scRNA-seq data. For this, we downloaded data from a recently published study of inflammatory skin conditions (Neuwirth, *et al*., 2025) and sought to recapitulate some of the results. Specifically, the first step of data integration and cell type identification and the last step of differential expression analysis between SAT1 low and SAT1 high expression regulatory T-cells. In both cases we corrected the data with reComBat-seq using the patient ID as the batch variable. Despite the complexity of scRNA-seq data, reComBat-seq managed to remove the batch variation well. Judging qualitatively by uniform manifold approximation and projection, we found good clustering by cell type where we previously had more clustering by patient ID (compare Figure 3A top and bottom; most notably the VE3 and F2 cluster). A similar situation was found when we compared uncorrected against corrected data of just regulatory T-cells (Figure 3B). To see if our observation in bulk differential expression analysis still holds up in scRNA-seq, we computed differentially expressed genes (DEGs) between health and disease regulatory T-cells using MAST and compared the recovered DEGs between uncorrected and corrected data. In total, we found 1,119 DEGs in the uncorrected data and 833 DEGs in the corrected data with an overlap of 826 DEGs (Figure 3C). Similarly, we observe a Spearman correlation coefficient of 0.977 between the two ranked result sets. Although, log2 fold changes (log2FC) are expectedly a bit lower in the corrected data compared to the uncorrected data, this result indicates a good agreement of log2 fold changes. In general, only lower log2FC genes are removed from DEGs after correction (Figure 3D). This recapitulates the results in bulk and again suggests a reduction in false positives by removing technical noise from the data. In summary, reComBat-seq is capable of handling the complex variance structure of scRNA-seq data. It effectively removes batch variation while again preserving biological signal.

**Figure 3:**
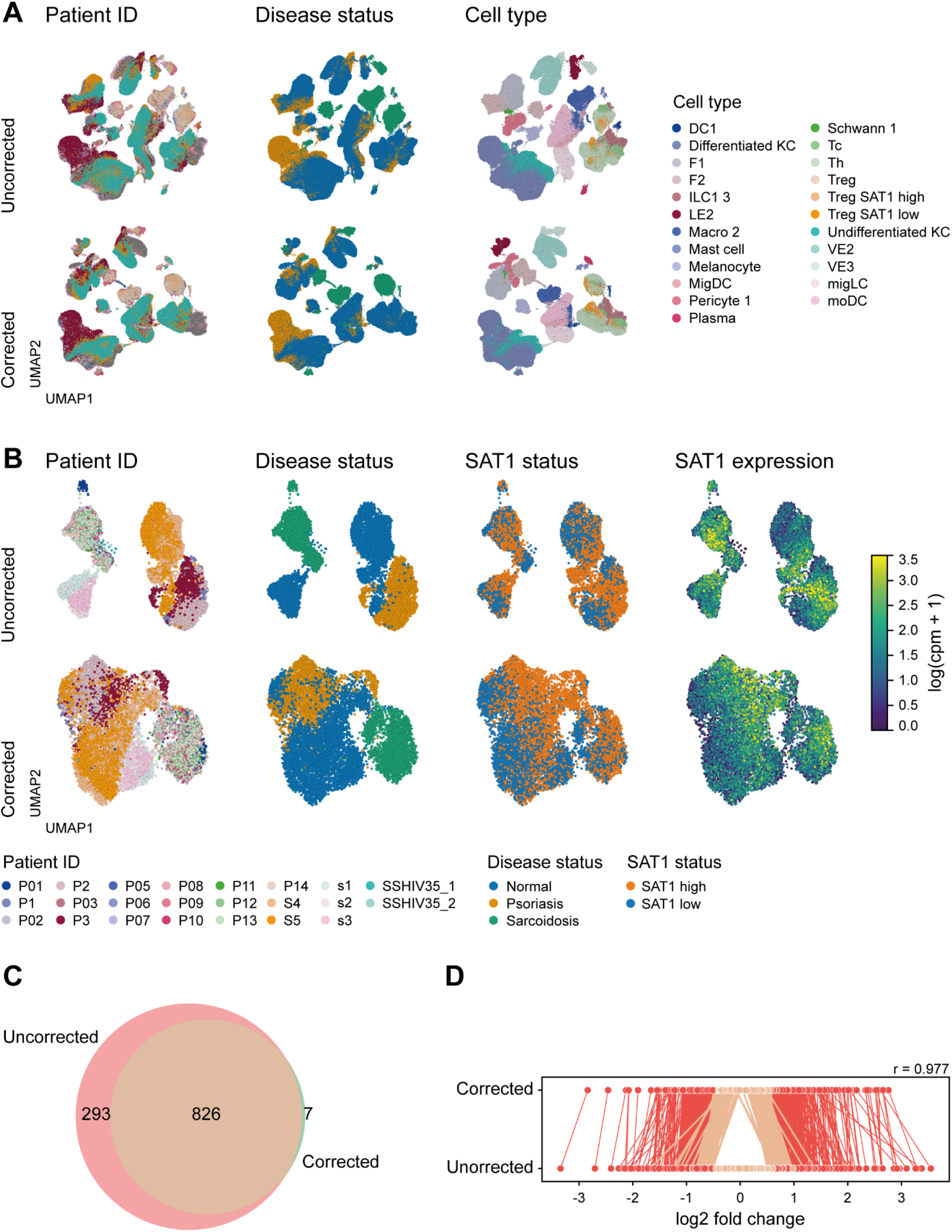
scRNA-seq data integration with reComBat-seq. (A) UMAPs of uncorrected and corrected data of 332,546 cells. Legends for Patient ID and Disease status below B. (B) UMAPs of regulatory T-cell subset. (C) Venn diagram of DEG overlap between uncorrected and corrected regulatory T-cell data. (D) Rank comparison of genes between uncorrected and corrected data based on log2 fold change. r indicates the Spearman correlation coefficient between the two lists of genes.

## Discussion

We developed reComBat-seq to close the gap between multi-experiment RNA-seq datasets and commonly used downstream analysis tools. Although parts of this have already been addressed by ComBat-seq (Zhang, *et al*., 2020), the introduced regularization now extends the methodology to even larger and possibly more complex datasets while still preserving the original nature of the data. We showed that reComBat-seq performs comparable to its methodological siblings while reducing false positives in downstream differential expression analyses. Furthermore, preserving counts provides a useful platform for better retention of biological signals after removal of technical noise which can be largely attributed to avoiding the aggressive data transformation into the Gaussian data distribution required by other comparable methods (Zhang, *et al*., 2020). This also enabled the use of state-of-the-art downstream analyses that require count data as input such as DESeq2 or edgeR. Here, especially the usage of DESeq2 showed that reComBat-seq stabilizes count variance and thus enables robust LFC estimation out of the box. We attribute this to a combination of the regularization and computing corrected counts directly from an estimated negative binomial distribution (Zhu, *et al*., 2019). Finally, reComBat-seq is also capable of removing considerable batch effects from single-cell RNA-seq data which often exhibits a complex source structure of batch effects (Luecken, *et al*., 2022). This makes reComBat-seq a suitable lightweight alternative to other more involved integration methods like scVI (Lopez, *et al*., 2018) or Harmony (Korsunsky, *et al*., 2019) again with the advantage of enabling typical downstream analyses by preserving the count nature of the data. While being applicable to most commonly used datasets, a remaining disadvantage of reComBat-seq is its current memory inefficiency, specifically with respect to sparse data such as scRNA-seq. The presented full scRNA-seq dataset consisted of 332,546 cells with 20,912 genes, demanding up to 831GB of RAM and 16h of runtime on 4 cores of an AMD EPYC 7662. This may be prohibitive for smaller compute clusters and is something to be improved upon in the future. In summary, we see reComBat-seq as a vital, count-native alternative for removing unwanted technical variation for the data-driven biology of today.

## Acknowledgements

We thank the Life Science Compute Cluster (LiSC) at the Centre for Microbiology and Environmental System Sciences of the University of Vienna for providing the computational resource necessary to perform the presented computations.

## References

Adamer, M.F. et al. (2022) reComBat: batch-effect removal in large-scale multi-source gene-expression data integration. Bioinformatics Advances, 2.

Anders, S. and Huber, W. (2010) Differential expression analysis for sequence count data. Genome Biol., 11, R106.

Chen, Y. et al. (2025) edgeR v4: powerful differential analysis of sequencing data with expanded functionality and improved support for small counts and larger datasets. Nucleic Acids Res., 53, gkaf018.

Di Meglio, P. et al. (2014) Activation of the aryl hydrocarbon receptor dampens the severity of inflammatory skin conditions. Immunity, 40, 989–1001.

Finak, G. et al. (2015) MAST: a flexible statistical framework for assessing transcriptional changes and characterizing heterogeneity in single-cell RNA sequencing data. Genome Biol., 16, 278.

Foulkes, A.C. et al. (2019) A framework for multi-omic prediction of treatment response to biologic therapy for psoriasis. J. Invest. Dermatol., 139, 100–107.

Goh, W.W.B. et al. (2022) Are batch effects still relevant in the age of big data? Trends Biotechnol., 40, 1029–1040.

Goh, W.W.B. et al. (2017) Why batch effects matter in omics data, and how to avoid them. Trends Biotechnol., 35, 498–507.

Gupta, R. et al. (2016) Landscape of long noncoding RNAs in psoriatic and healthy skin. J. Invest. Dermatol., 136, 603–609.

Himes, B.E. et al. (2014) RNA-Seq transcriptome profiling identifies CRISPLD2 as a glucocorticoid responsive gene that modulates cytokine function in airway smooth muscle cells. PLoS One, 9, e99625.

Hui, H.W.H. et al. (2024) Thinking points for effective batch correction on biomedical data. Brief. Bioinform., 25, bbae515.

Johnson, W.E. et al. (2007) Adjusting batch effects in microarray expression data using empirical Bayes methods. Biostatistics, 8, 118–127.

Korsunsky, I. et al. (2019) Fast, sensitive and accurate integration of single-cell data with Harmony. Nat. Methods, 16, 1289–1296.

Leek, J.T. et al. (2010) Tackling the widespread and critical impact of batch effects in high-throughput data. Nat. Rev. Genet., 11, 733–739.

Li, B. et al. (2014) Transcriptome analysis of psoriasis in a large case-control sample: RNA-seq provides insights into disease mechanisms. J. Invest. Dermatol., 134, 1828–1838.

Lopez, R. et al. (2018) Deep generative modeling for single-cell transcriptomics. Nat. Methods, 15, 1053–1058.

Love, M.I. et al. (2014) Moderated estimation of fold change and dispersion for RNA-seq data with DESeq2. Genome Biol., 15, 550–550.

Luecken, M.D. et al. (2022) Benchmarking atlas-level data integration in single-cell genomics. Nat. Methods, 19, 41–50.

Luecken, M.D. and Theis, F.J. (2019) Current best practices in single-cell RNA-seq analysis: a tutorial. Mol. Syst. Biol., 15, e8746.

Malzl, D. and Neuwirth, T. (2024) processed scRNA-seq data for Neuwirth & Malzl et al. 2024.

Neuwirth, T. et al. (2025) The polyamine-regulating enzyme SSAT1 impairs tissue regulatory T cell function in chronic cutaneous inflammation. Immunity, 58, 632–647.e12.

Patel, H., Beber, M.E., et al. (2022) nf-core/fetchngs: nf-core/fetchngs v1.7 - Sodium Skunk Zenodo.

Patel, H., Ewels, P., et al. (2022) nf-core/rnaseq: nf-core/rnaseq v3.7 - Iron Iguana Zenodo.

Ritchie, M.E. et al. (2015) limma powers differential expression analyses for RNA-sequencing and microarray studies. Nucleic Acids Res., 43, e47.

Tran, H.T.N. et al. (2020) A benchmark of batch-effect correction methods for single-cell RNA sequencing data. Genome Biol., 21, 12.

Wolf, F.A. et al. (2018) SCANPY: large-scale single-cell gene expression data analysis. Genome Biol., 19, 15.

Yu, Y. et al. (2024) Assessing and mitigating batch effects in large-scale omics studies. Genome Biol., 25, 254.

Yu, Z. et al. (2020) High-throughput transcriptome and pathogenesis analysis of clinical psoriasis. J. Dermatol. Sci., 98, 109–118.

Zappia, L. et al. (2017) Splatter: simulation of single-cell RNA sequencing data. Genome Biol., 18, 174.

Zhang, X. (2025) Highly effective batch effect correction method for RNA-seq count data. Comput. Struct. Biotechnol. J., 27, 58–64.

Zhang, Y. et al. (2020) ComBat-seq: batch effect adjustment for RNA-seq count data. NAR Genom. Bioinform., 2, qaa078.

Zhu, A. et al. (2019) Heavy-tailed prior distributions for sequence count data: removing the noise and preserving large differences. Bioinformatics, 35, 2084–2092.

