## supplementary file for "reComBat-seq: Regularized negative binomial regression for batch-effect correction in underdetermined transcriptomics datasets"

### Supplementary Figures

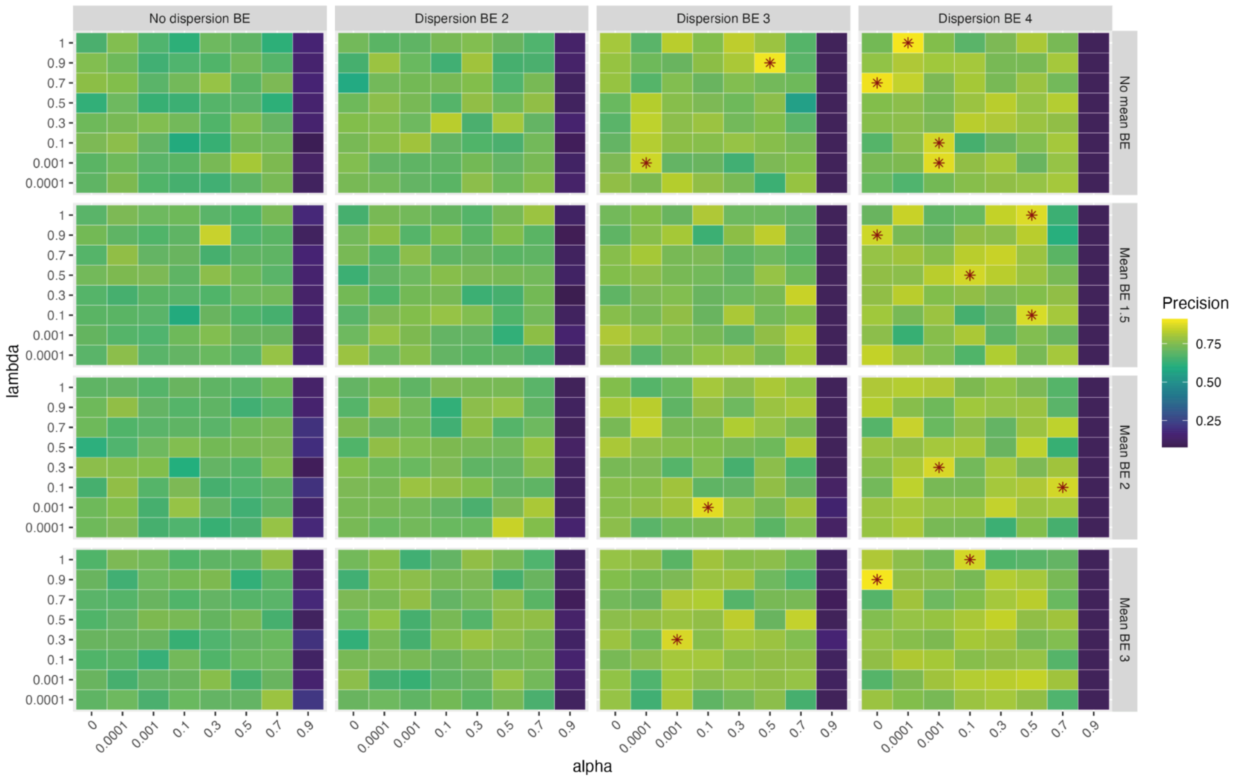


**Figure S1:** Overview over precision score across different regularization parameters and batch fold effects. The heat-map showcases mean values for the precision over 5 repeated runs evaluated on synthetic data (128 samples, 64 samples/batch, 2 biological groups). Parameter settings that yielded a mean precision score higher than 0.8 were marked with a star. Values of $\alpha > 0.9$ led to unstable behavior, resulting in extremely low precision scores (as low as 0.1) or, in some cases, infinite loops that required manual interruption of the algorithm.

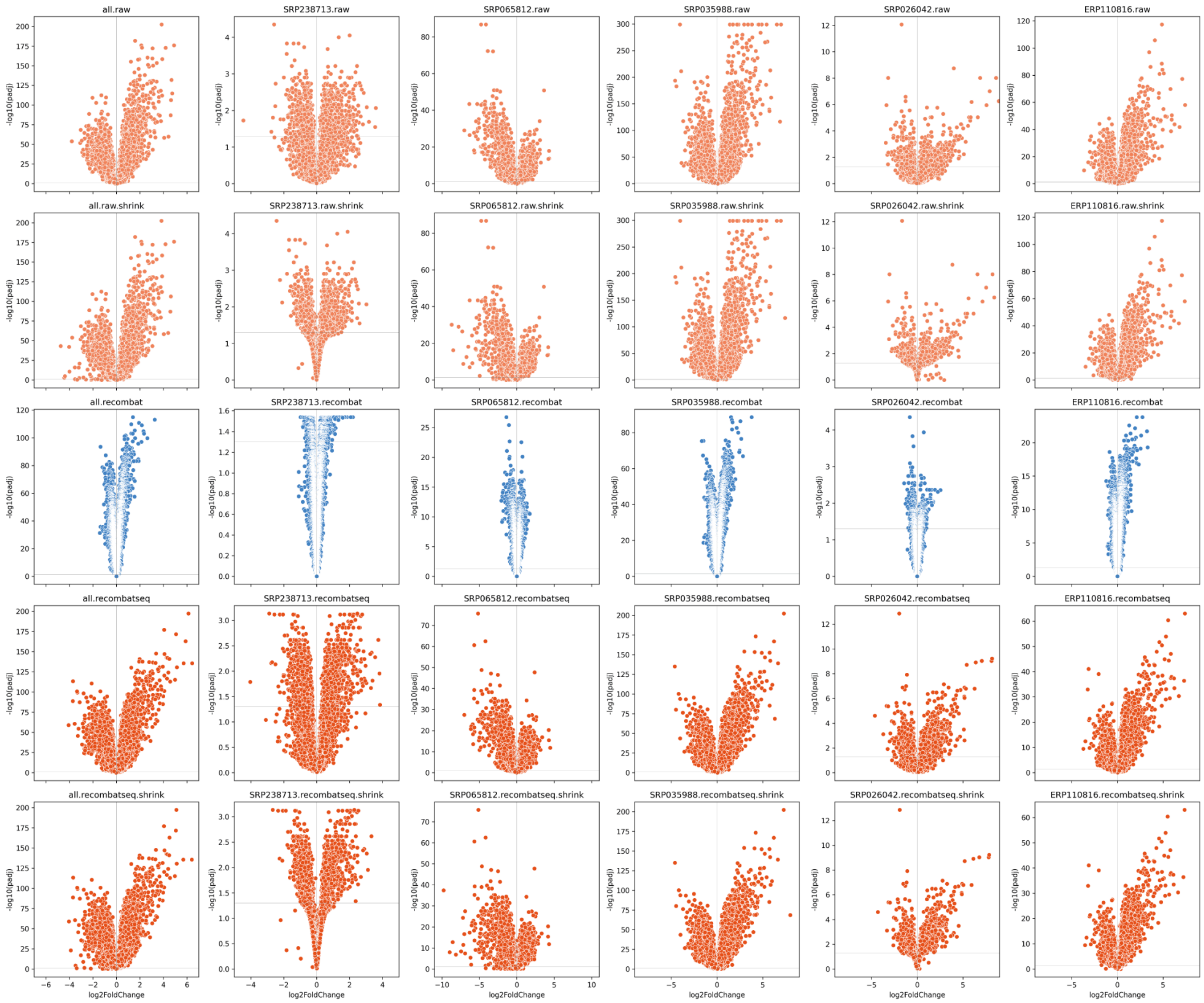


**Figure S2:** Volcano plots for bulk RNA-seq differential expression analysis of full and individual uncorrected, corrected data. Horizontal lines mark padj < 0.05 threshold.

### From ComBat-seq to reComBat-seq: Derivation of regularized negative binomial regression

ComBat-seq utilizes a generalized linear model (GLM) to capture the discrete, overdispersed nature of RNA-seq count data. It assumes the counts follow a negative binomial distribution $Y_{ijg} \sim NB\left(\mu_{ijg}, \phi_{ig}\right)$, with a log-link function connecting the mean to the linear predictor:

$$\log\ \mu_{ijg} = (X\beta)_{jg} + \log N_j.$$

The probability mass function is parameterised by:

$$f\left(y_{jg}; \mu_{jg}, \phi_g\right) = \frac{\Gamma\left(y_{jg}+\frac{1}{\phi_g}\right)}{\Gamma\left(\frac{1}{\phi_g}\right)\Gamma(y+1)}\left(\frac{1}{1+\phi_g\mu_{jg}}\right)^{\left(\frac{1}{\phi_g}\right)}\left(\frac{\mu_{jg}}{1+\phi_g\mu_{jg}}\right)^{y}.$$

The regression coefficients $\beta_g = \left(\beta_{g1}, \ldots, \beta_{gp}\right)$ are typically estimated using maximum likelihood methods (MLE) by minimising the negative log-likelihood function. The likelihood function is calculated using the probability mass function: $L\left(\mu_{jg}, \phi_g\right) = \prod_{j=1}^{n} f(y_{jg}; \mu_{jg}, \phi_g)$.

Taking the logarithm gives us the (negative) log-likelihood function:

$$\begin{aligned} NLL\left(\beta_g, \phi_g\right) &= -LL\left(\beta_g, \phi_g\right) = -\log L\left(\mu_{jg}, \phi_g\right) \\ &= -\sum_{j=1}^{n} \log\Gamma\left(y_{jg} + \frac{1}{\phi_g}\right) - \log\Gamma\left(y_{jg} + 1\right) - \log\Gamma\left(\frac{1}{\phi_g}\right) + \\ &\quad y_{jg}\log\phi_g + y_{jg}\log\mu_{jg} - \left(y_{jg} + \frac{1}{\phi_g}\right)\log\left(1 + \phi_g\mu_{jg}\right) \end{aligned}$$

The sample specific row of the design matrix is denoted by $x_j = \left(x_{j1}, \ldots, x_{jn}\right)$. The gradient of the link function $\mu_{jg} = e^{x_j\beta_g}$ is calculated as: $\frac{\partial\mu_{jg}}{\partial\beta_{gk}} = \frac{\partial e^{x_j\beta_g}}{\partial\beta_{gk}} = x_{jk}e^{x_j\beta_g} = x_{jk}\mu_{jg}$. To estimate $\beta_g$ we find the roots of the score equations $\frac{\partial LL\left(\beta_g, \phi_g\right)}{\partial\beta_{gk}} = 0$. Differentiating with respect to the $k$-th coefficient $\beta_{gk}$:

$$\begin{aligned} \frac{\partial LL\left(\beta_g, \phi_g\right)}{\partial\beta_{gk}} &= \frac{\partial}{\partial\beta_{gk}} \sum_{j=1}^{n} y_{jg}\ \log\mu_{jg} - \left(y_{jg} + \frac{1}{\phi_g}\right)\log\left(1 + \phi_g\mu_{jg}\right) \\ &= \sum_{j=1}^{n} y_{jg}x_{jk}\mu_{jg}\frac{1}{\mu_{jg}} - \left(y_{jg} + \frac{1}{\phi_g}\right)\frac{1}{1+\phi_g\mu_{jg}}\left(\phi_g x_{jk}\mu_{jg}\right) \end{aligned}$$

$$= \sum_{j=1}^{n} \frac{x_{jk}\left(y_{jg} - \mu_{jg}\right)}{1+\phi_g \mu_{jg}}$$

As there is often no closed-form solution for $\beta_g$, we employ the Fisher scoring algorithm. It replaces the observed Hessian $(H)$ with its expected value, the Fisher information matrix $(I(\beta))$, defined as:

$$I(\beta) \;=\; E[-\;H] = E\left[-\frac{\partial^2 LL(\beta, \phi)}{\partial\beta^2}\right] = E\left[\frac{\partial^2 NLL(\beta, \phi)}{\partial\beta^2}\right]$$

The iterative update step is defined as:

$$\beta^{(n+1)} \;=\beta^{(n)} - \left(I\left(\beta^{(n)}\right)\right)^{-1}\left(\frac{\partial NLL\left(\beta^{(n)}, \phi\right)}{\partial\beta^{(n)}}\right) \;=\; \beta^{(n)} + \left(I\left(\beta^{(n)}\right)\right)^{-1}\left(\frac{\partial LL\left(\beta^{(n)}, \phi\right)}{\partial\beta^{(n)}}\right)$$

The entries of the Fisher information matrix are derived from the second derivatives of the log-likelihood function:

$$H_{lk} \;=\; \frac{\partial^2 NLL}{\partial\beta_l \partial\beta_k} \;=\; \frac{\partial NLL}{\partial\beta_l}\sum_{j=1}^{n}\frac{x_{jk}\left(y_{jg}-\mu_{jg}\right)}{1+\phi_g\mu_{jg}} \;=\; \sum_{j=1}^{n} x_{jk}\frac{-x_{jl}\mu_j\left(1+\phi_g\mu_{jg}\right) - \left(y_{jg}-\mu_{jg}\right)\phi_g x_{jl}\mu_{jg}}{\left(1+\phi_g\mu_{jg}\right)^2}$$

$$I\left(\beta_g\right)_{lk} \;=\; E\left[-\;H_{lk}\right] \;=\; -\sum_{j=1}^{n} x_{jk}\left(E\left[\frac{-x_{jl}\mu_j\left(1+\phi_g\mu_{jg}\right)}{\left(1+\phi_g\mu_{jg}\right)^2}\right] - E\left[\frac{\left(y_{jg}-\mu_{jg}\right)\phi_g x_{jl}\mu_{jg}}{\left(1+\phi_g\mu_{jg}\right)^2}\right]\right) \;=\; \sum_{j=1}^{n} x_{jk}x_{jl}\frac{\mu_{jg}}{1+\phi_g\mu_{jg}}$$

Defining a diagonal weight matrix $W^n$ with elements $w_{jj}^{\;\;n} \;=\; \frac{\mu^n_{\;jg}}{1+\phi_g\mu^n_{\;jg}}$, the information matrix can be rewritten in a compact form:

$$I(\beta^{(n)})_{lk} \;=\sum_{j=1}^{n} x_{jk}x_{jl}\frac{\mu^{(n)}_{\;jg}}{1+\phi_g\mu^{(n)}_{\;\;jg}} \;=\left(X^T W^{(n)} X\right)_{lk}$$

By introducing the adjusted dependent variable $z^{(n)}_{\;\;j} \;=\; x_j\beta^{(n)} \;+\; \frac{y_j - \mu^{(n)}_{\;\;j}}{\mu^{(n)}_{\;\;j}}$, the update simplifies to the Iteratively reweighted least squares (IRLS) form:

$$I(\beta^{(n)})\beta^{(n+1)} \;=\; I(\beta^{(n)})\beta^{(n)} \;+\; \left(\frac{\partial LL\left(\beta^{(n)}_{\;g}, \phi_g\right)}{\partial\beta^{(n)}_{\;\;gk}}\right)_k$$

$$(X^T W^{(n)} X)\beta^{(n+1)} \;= (X^T W^{(n)} X)\beta^{(n)} \;+\left(\sum_{j=1}^{n}\frac{x_{jk}\left(y_{jg}-\mu^{(n)}_{\;\;jg}\right)}{1+\phi_g\mu^{(n)}_{\;\;jg}}\right)_k$$

$$(X^T W^{(n)} X)\beta^{(n+1)} = (X^T W^{(n)} X)\beta^{(n)} + X^T W^{(n)} \left( \frac{y_j - \mu^{(n)}_{jg}}{\mu_{jg}} \right)_j$$

$$\beta^{(n+1)} = (X^T W^{(n)} X)^{-1} X^T W^{(n)} \left( X\beta^{(n)} + \left( \frac{y_j - \mu^{(n)}_{jg}}{\mu^{(n)}_{jg}} \right)_j \right)$$

$$\beta^{(n+1)} = (X^T W^{(n)} X)^{-1} X^T W^{(n)} z^{(n)}$$

The algorithm relies on finding the inverse of the Fisher information matrix to estimate the parameters. If $X$ is singular, then $X^T WX$ will also be singular and thus non-invertible. reComBat-seq introduces elastic net penalties to ensure the strict convexity and stability of the system:

$$NLL_{reg} = NLL\left(\beta_g, \phi_g\right) + \lambda\left(\alpha\, ||\beta||_1 + \left(\frac{1-\alpha}{2}\right) ||\beta||_2^2\right)$$

The penalized Fisher information matrix becomes:

$$I(\beta^{(n)})_{reg} = X^T W^{(n)} X + \lambda(1 - \alpha)I$$

and the gradient of $NLL_{reg}$ is adjusted as:

$$\left( \frac{\partial NLL_{reg}\left(\beta^{(n)}; y\right)}{\partial \beta} \right)_k = \left( \frac{\partial NLL_{reg}\left(\beta^{(n)}; y\right)}{\partial \beta} \right)_k + \lambda\left(\alpha\, sgn(\beta_k) + (1 - \alpha)\beta_k\right).$$
